# ElectroEncephaloGraphy robust statistical linear modelling using a single weight per trial

**DOI:** 10.1101/2021.04.27.441629

**Authors:** Cyril Pernet, Guillaume Rousselet, Ignacio Suay Mas, Ramon Martinez, Rand Wilcox, Arnaud Delorme

## Abstract

Being able to remove or weigh down the influence of outlier data is desirable for any statistical models. While Magnetic and ElectroEncephaloGraphic (MEEG) data are often averaged across trials per condition, it is becoming common practice to use information from all trials to build statistical linear models. Individual trials can, however, have considerable weight and thus bias inferential results (effect sizes as well as thresholded t/F/p maps). Here, rather than looking for univariate outliers, defined independently at each measurement point, we apply the principal component projection (PCP) method at each channel, deriving a single weight per trial at each channel independently. Using both synthetic data and open EEG data, we show (1) that PCP is efficient at detecting a large variety of outlying trials; (2) how PCP-based weights can be implemented in the context of the general linear model (GLM) with accurate control of type 1 family-wise error rate; and (3) that our PCP-based Weighted Least Square (WLS) approach increases the statistical power of group analyses as well as a much slower Iterative Reweighted Least Squares (IRLS), although the weighting scheme is markedly different. Together, our results show that WLS based on PCP weights derived from whole trial profiles is an efficient method to weigh down the influence of outlier EEG data in linear models.

**Data availability:** all data used are publicly available (CC0), all code (simulations and data analyzes) is also available online in the LIMO MEEG GitHub repository (MIT license).

## Introduction

MEEG data analysis can be subdivided into two main parts: preprocessing and processing. Data preprocessing corresponds to any manipulation and transformation of the data, such as artefacts removal and attenuation or change in data representations (spectral transformation, source modelling). Data processing refers to mathematical procedures that do not change the data, i.e., statistical analysis and statistical modeling (1). Here, we present a method that improves data processing by weighting down trials which have different dynamics from the bulk of the data, within the context of an experimental design. We investigated single trial weighting for full scalp/channel statistical linear hierarchical modelling, providing a more robust statistical analysis method to the community.

After data preprocessing, MEEG data are typically epoched to form 3 or 4-dimensional matrices of, e.g., channel x time x trials and channel x frequency x time x trials. Several neuroimaging packages are dedicated to the statistical analyses of such large multidimensional data, often using linear methods. For instance, in the LIMO MEEG toolbox (2), each channel, frequency, and time frame is analyzed independently using the general linear model, an approach referred to as mass-univariate analysis. Ordinary Least Squares (OLS) are used to find model parameters that minimize the error between the model and the data. For least squares estimates to have good statistical properties, it is however expected that the error covariance off-diagonals are zeros, such that Cov(e) = σ^2^I, I being the identity matrix (3), assuming observations are independent and identically distributed. It is well established that deviations from that assumption lead to substantial power reduction and to an increase in the false-positive rate. When OLS assumptions are violated, robust techniques offer reliable solutions to restore power and control the false positive rate. Weighted Least Squares (WLS) is one such robust method that uses different weights across trials, such that Cov(e) = σ^2^V, with V a diagonal matrix:

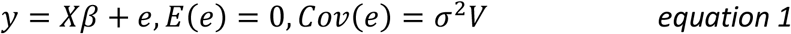

with *y* a n-dimensional vector (number of trials), *X* the n*p design matrix, *β* a p dimensional vector (number of predictors in *X*) and *e* the error vector of dimension n. The WLS estimators can then be obtained using an OLS on transformed data (eq. 2 and 3):

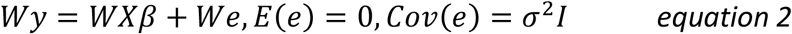

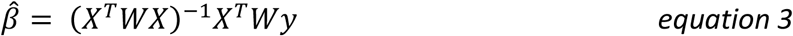

with *W*a 1*n vector of weights.

When applied to MEEG data, a standard mass-univariate WLS entails obtaining a weight for each trial but also each dimension analyzed, i.e. channels, frequencies and time frames. Following such procedure, a trial could be considered as an outlier or be assigned a low weight, for a single frequency or time frame, which is implausible given the well-known correlations of MEEG data over space, frequencies and time. We propose here that a single or a few consecutive data points should never be flagged as outliers or weighted down, and that a single weight per trial (and channel) should be derived instead, with weights taking into account the whole temporal or spectral profile. In the following, we demonstrate how the Principal Component Projection method (PCP (4)) can be used in this context, and how those weights can then be used in the context of the general linear model (WLS), applied here to event-related potentials. Such weighting should not be taken to bypass data preprocessing. Measurements and physiological artefacts are typically several order of magnitude stronger than the signal of interest, spanning consecutive trials, and require dedicated algorithms. After data preprocessing, residual signal artefacts might however remain. In addition, natural and unintended experimental variations might also exist among all trials, and the weighting scheme aims to account for these.

## Method

### Trial-based Weighted Least Squares

The new WLS solution consists of computing 1^st^ level GLM beta estimates using weights from the Principal Component Projection algorithm (PCP - (4)). As such, the estimation procedure follows the usual steps of weighting schemes, like a standard Iterative Reweighted Least Squares (IRLS) solution:

1. After the OLS solution is computed, an adjustment is performed on residuals by multiplying them by 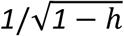 where h is a vector of Leverage points (i.e. the diagonal of the hat matrix *H*=*X*(*X*^′^*X*)^−*1*^*X*^′^ where X is the design matrix reflecting experimental conditions). This adjustment is necessary because leverage points are the most influential on the regression space, i.e. they tend to have low residual values (5).
2. Residuals are then standardized using a robust estimator of dispersion, the median absolute deviation to the median (MAD), and re-adjusted by the tuning function. Here we used the bisquare function. A series of weights is next obtained either based on distances (PCP method) or minimizing the error term (IRLS) with, in both cases, high weights for data points having high residuals (with a correction for Leverage).
3. The solution is then computed following equation 3.

Unlike IRLS, WLS weights are not derived for each time frames independently but by looking at the multivariate distances in the principal component space (step 2 above) and thus deriving a unique weight for all time frames. An illustration of the method is shown in figure 1. Trial weights are computed as a distance among trials projected onto the main (>=99%) principal components space. Here, the principal components computed over the f time frames are those directions which maximize the variance across trials for uncorrelated (orthogonal) time periods (figure 1B). Outlier trials are points in the f-dimensional space which are far away from the bulk. By virtue of the PCA, these outlier trials become more visible along the principal component axes than in the original data space. Weights (figure 1E) for each trial are obtained using both the Euclidean norm (figure 1C, distance location) and the kurtosis weighted Euclidean norm (figure 1D, distance scatter) in this reduced PCA space (see (4) for details). Weights are then applied to each trial (at each data points), on the original data (i.e. there is no data reduction). We exploit this simple technique because it is computationally fast given the rich dimensional space of EEG data and because it does not assume the data to originate from a particular distribution. The PCP algorithm is implemented in the *limo_pcout*.*m* function, and the WLS solution in the *limo_WLS*.*m* function, both distributed with the LIMO MEEG toolbox (https://limo-eeg-toolbox.github.io/limo_meeg/).

**Figure 1.**
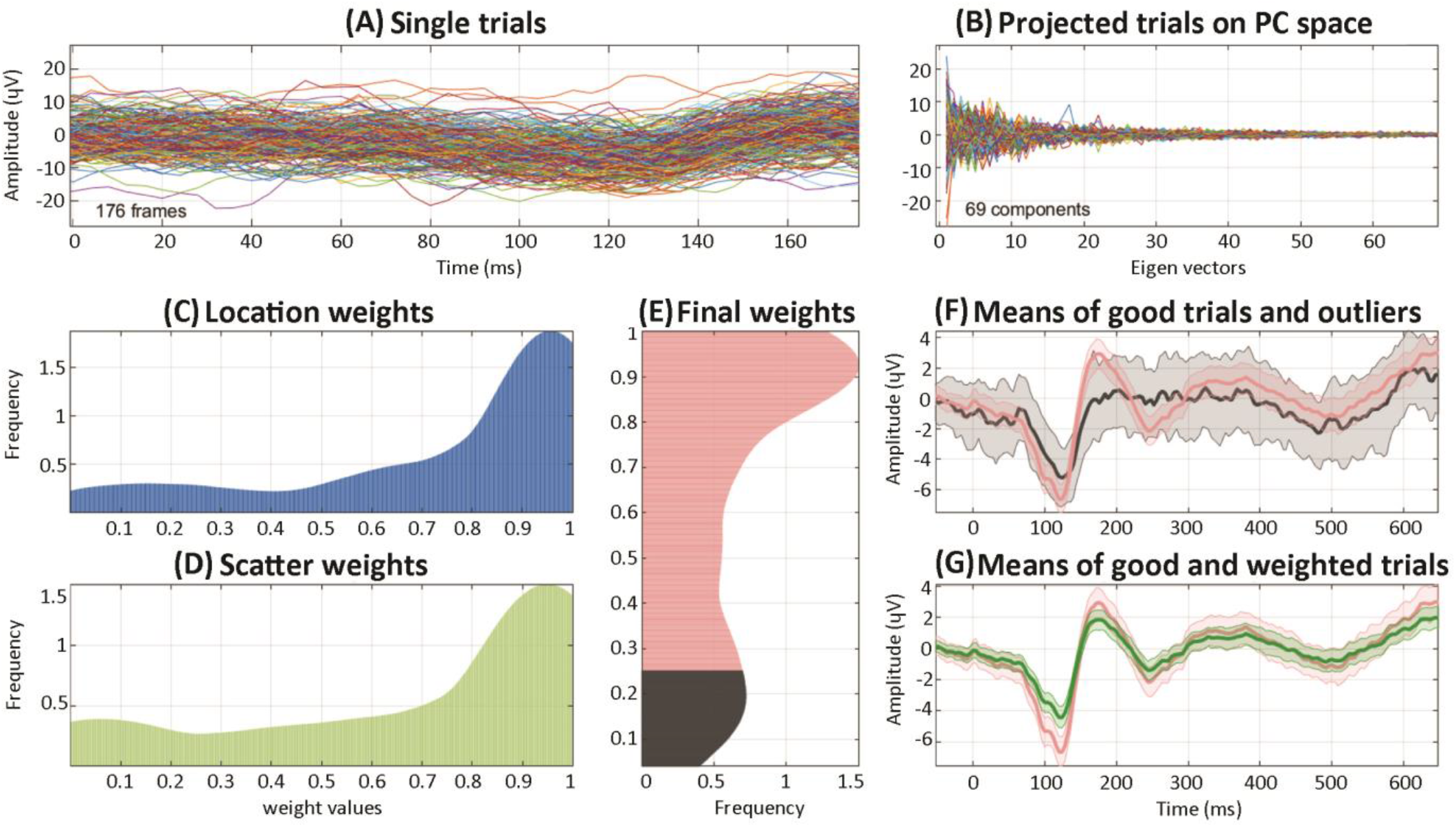
Illustration of the PCP weighting scheme using trials for ‘famous faces’ of the OpenNeuro.org publicly available ds002718 dataset. Data are from subject 3, channel 34 (see Section on empirical data analysis). Panel A shows the single-trial responses to all stimuli. The principal component analysis is computed over time, keeping the components explaining the most variance and summing to at least 99% of explained variance (giving here 69 eigenvectors i.e. independent time components from the initial 176 time points). The data are then projected onto those axes (panel B). From the data projected onto the components, Euclidean distances for location and scatter are computed (panels C, D - showing smooth histograms of weights) and combined to obtain a distance for each trial. That distance is either used as weights in a linear model or used to determine outliers (panel E, with outliers identified for weights below ∼0.27, shown in dark grey). At the bottom right, the mean ERP for trials classified as good (red) vs. outliers (black) and the weighted mean (green) are shown (panels F and G). Shaded areas indicate the 95% highest-density percentile bootstrap intervals.

### Simulation-based analyses

#### A. Outlier detection and effectiveness of the weighting scheme

Simulated ERPs were generated to evaluate the classification accuracy of the PCP method and to estimate the robustness to outliers and signal-to-noise ratio of the WLS solution in comparison to an OLS solution and a standard Iterative Reweighted Least Squares (IRLS) solution, which minimizes residuals at each time frame separately (implemented in *limo_IRLS*.*m*). To do so, we manipulated (i) the percentage of outliers, using 10%, 20%, 30%, 40% or 50% of outliers; (ii) the signal to noise ratio (defined relative to the mean over time of the background activity); and (iii) the type of outliers. The first set of outliers were defined based on the added noise: white noise, pink noise, alpha oscillations and gamma oscillations following (6). In these cases, the noise started with a P1 component and lasted ∼ 200ms (see below). The second set of outliers were defined based on their amplitude, or outlier to signal ratio (0.5, 0.8, 1.2, and 1.5 times the true N1 amplitude).

Synthetic data were generated for one channel, using the model developed by Yeung et al. (2018 - (7)). The simulated signal corresponded to an event-related potential with P1 and N1 components (100 ms long) added to background activity with the same power spectrum as human EEG, generating 200 trials of 500 ms duration with a 250 Hz sampling rate. Examples for each type of simulation are shown in figure 2 and results are based, for each case, on a thousand random repetitions. Performance of the PCP algorithm at detecting outlying synthetic EEG trials was investigated by computing the confusion matrix and mapping the true and false positives rates in the Receiver Operating space, and by computing the Matthew Correlation Coefficients (MCC – equation 4). The robustness, or effectiveness of the weighting scheme, was examined by computing the Pearson correlations and the Kolmorov-Smirnov (KS) distances between the ground truth mean and the OLS, WLS, and IRLS means. Pearson values allowed to estimate the linear relationships between estimated means and the truth while KS distances provide a fuller picture of the overall differences in distributions.

**Figure 2.**
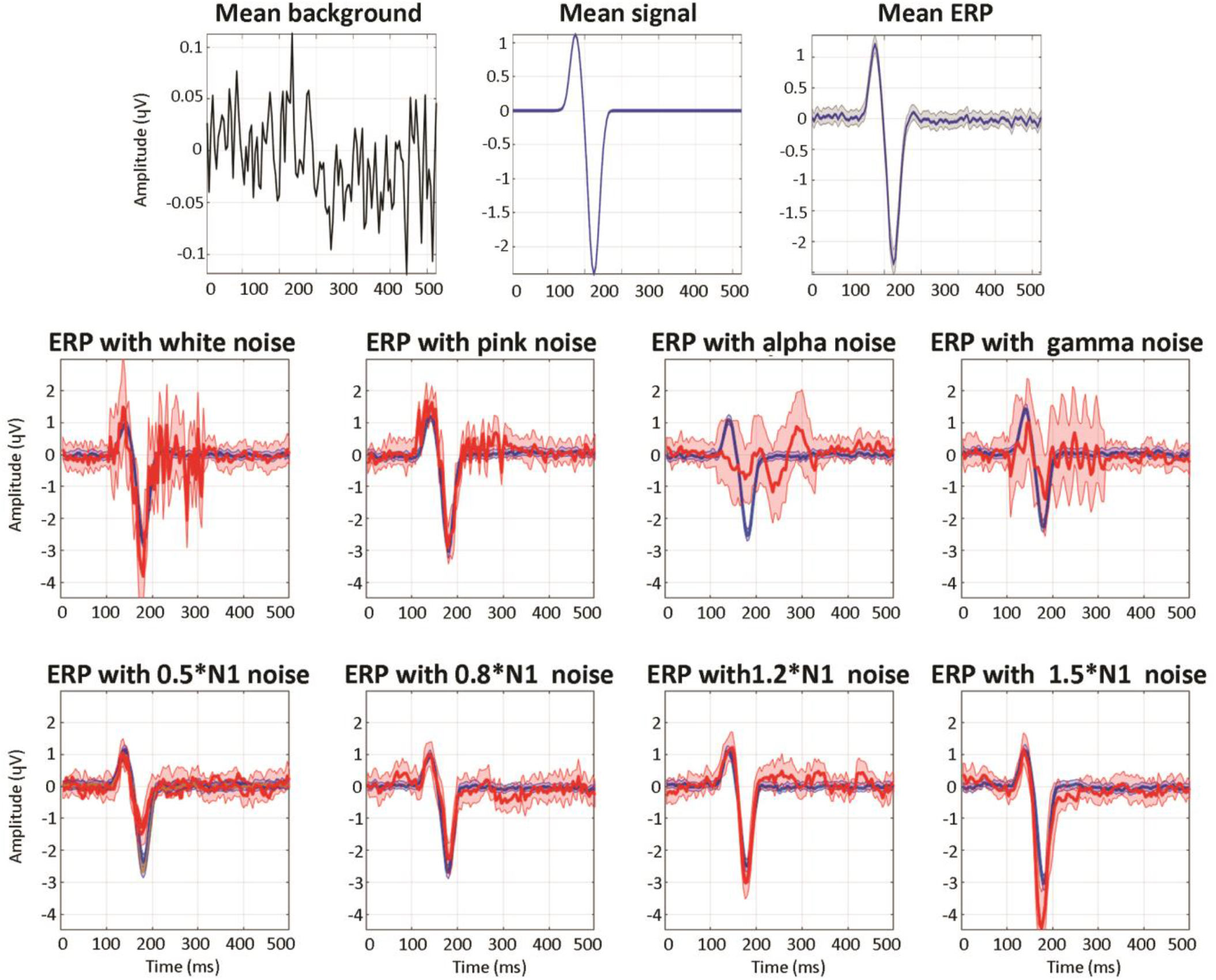
Illustration of simulated ERP ground truth with the different types of outlier trials. At the top is shown the mean background, mean signal and resulting generated ERP with it’s 95% confidence intervals. In each subsequent subplot is shown the mean ERP ground truth from 160 trials with their 95% confidence intervals (blue) with a SNR of 1. The first row shows in red the mean ERP from outlier trials generated by adding white noise, pink noise, alpha or gamma oscillations; the second row shows the mean ERP from outlier trials generated with variable Outlier to Signal Ratio (OSR) on the N1 component.

The code used to generate the ERP and the results are available at https://github.com/LIMO-EEG-Toolbox/limo_test_stats/tree/master/PCP_simulations/.

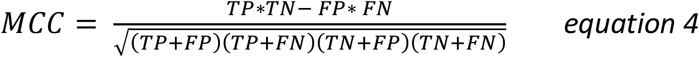

with TP the true positives, TN the true negatives, FP the false positives and FN the false negatives.

#### B. Statistical inference

Accurate estimation of model parameters (i.e. beta estimates in the GLM - equation 3) is particularly important because it impacts group-level results. Inference at the single-subject level may, however, also be performed and accurate p-values need to be derived. Here, error degrees of freedom are obtained using the Satterthwaite approximation ((8) equation 5).

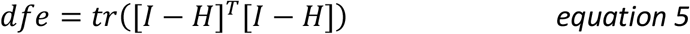

### with dfe, the degree of freedom of the error, I the identity matrix, and H the hat matrix

To validate p-values, simulations under the null were performed. Two types of data were generated: Gaussian data of size 120 trials x 100 time frames and EEG data of size 120 trials x 100 time frames with a P1 and N1 component as above, added to coloured background activity with the same power spectrum as human EEG. In each case, a regression (1 Gaussian random variable), an ANOVA (3 conditions of 40 trials - dummy coding) and an ANCOVA (3 conditions of 40 trials and 1 Gaussian random covariate) model were fitted to the data using the OLS, WLS and IRLS methods. The procedure was performed 10,000 times, leading to 1 million p-values per data/model/method combination and Type 1 errors with binomial confidence intervals were computed.

### Empirical data analysis

A second set of analyses used the publicly available multimodal face dataset (9) to (i) investigate the PCP classification; (ii) validate the GLM implementation for type 1 error family-wise control at the subject level; (iii) evaluate group results, contrasting WLS against the OLS and IRLS methods. This analysis can be reproduced using the script available at https://github.com/LIMO-EEG-Toolbox/limo_meeg/blob/master/resources/code/Method_validation.m.

#### A. EEG Data and Preprocessing

The experiment consisted in the presentation of familiar, unfamiliar, and scrambled faces, repeated twice at various intervals, leading to a factorial 3 (type of faces) by 3 (repetition) design. The preprocessing reused the pipeline described in Pernet et al (2021)(10). EEG data were extracted from the MEG fif files, time corrected and electrode position re-oriented and saved according to EEG-BIDS (11) (available at <u>OpenNeuro 10.18112/openneuro.ds002718.v1.0.2</u>.). Data were imported into EEGLAB (12) using *the bids-matlab-tools v5*.*2 plug-in* and non-EEG channel types were removed. Bad channels were next automatically removed and data filtered at 0.5 Hz using *pop_clean_rawdata*.*m* of the *clean_radata* plugin v2.2 (transition band [0.25 0.75], bad channel defined as a flat line of at least 5 sec and with a correlation to their robust estimate based on other channels below 0.8). Data were then re-referenced to the average (*pop_reref*.*m*) and submitted to an independent component analysis (13) (*pop_runica*.*m* using the runnica algorithm sphering data by the number of channels -1). Each component was automatically labelled using the *ICLabel* v1.2.6 plug-in (14), rejecting components labeled as eye movements and muscle activity above 80% probability. Epochs were further cleaned if their power deviated too much from the rest of the data using the Artifact Subspace Reconstruction algorithm (15) (*pop_clean_rawdata*.*m*, burst criterion set to 20).

#### B. High vs. low weight trials and parameters estimation

At the subject level (1st level), ERP were modelled at each channel and time frame with the 9 conditions (type of faces x repetition) and beta parameter estimates obtained using OLS, WLS, and IRLS. For each subject, high vs. low weight trials were compared with each other at the channel showing the highest between trials variance to investigate what ERP features drove the weighting schemes. High and low trials were defined a priori as trials with weights (or mean weights for IRLS) below the first decile or above the 9th decile. We used a two-sample bootstrap-t method to compare the 20% trimmed means of high and low trials in every participant, for each of these three quantities: temporal SNR (the standard deviation over time); global power (mean of squared absolute values, Parseval’s theorem); autocorrelation (distance between the 2 first peaks of the power spectrum density, Wiener-Khinchin theorem). A similar analysis was conducted at the group level averaging the metrics across trials. Computations of the three quantities have been automatized for LIMO MEEG v3.0 in the *limo_trialmetric*.*m* function.

#### C. Statistical inference

In mass-univariate analyses, once p-values are obtained, the family-wise type 1 error rate can be controlled using the distribution of maxima statistics from data generated under the null hypothesis (16). Here, null distributions were obtained by first centering data per conditions, i.e. the mean is subtracted from the trials in each condition, such that these distributions had a mean of zero, but the shape of the distributions is unaffected. We then bootstrap these centred distributions (by sampling with the replacement), keeping constant the weights (since they are variance stabilizers) and the design. We computed 2500 bootstrap estimates per subject. A thousand of these bootstrap estimates were used to compute the family-wise type 1 error rate (FWER), while maxima and cluster maxima distributions were estimated using from 300 to 1,500 bootstraps estimates in steps of 300, to determine the convergence rate, i.e. the number of resamples needed to control the FWER. Since OLS was already validated in Pernet et al. (2015), here we only present WLS results. Statistical validations presented here and other statistical tests implemented in the LIMO MEG toolbox v3.0 (GLM validation, robust tests, etc.) are all available at https://github.com/LIMO-EEG-Toolbox/limo_test_stats/wiki.

#### D. Performance evaluation at the group level

At the group level (2nd level), we computed 3 by 3 repeated measures ANOVAs (Hotelling T^^2^ tests) separately on OLS, WLS, and IRLS estimates, with the type of faces and repetition as factors. Results are reported using both a correction for multiple comparisons with cluster-mass and with TFCE (threshold-free cluster enhancement) at p<.05 (16,17).

In addition to these thresholded maps, distributions were compared to further understand where differences originated from. First, we compared raw effect sizes (Hotelling T^2) median differences between WLS vs. OLS and WLS vs. IRLS for each effect (face, repetition and interaction), using a percentile t-test with alpha adjusted across all 6 tests using Hochberg’s step-up procedure (18). This allowed checking if differences in results were due to effect size differences. Then, since multiple comparison correction methods are driven by the data structure, we compared the shapes of the F value and of the TFCE value distributions (TFCE reflecting clustering). Each distribution was standardized (equation 6) and WLS vs. OLS and WLS vs. IRLS distributions compared at multiple quantiles (19).

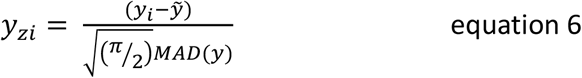

*with y*_*zi*_ *the standardized data, y the data*, 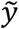 *the median and MAD the median absolute deviation*.

## Results

### Outlier detection

The PCP method is used in the GLM to obtain weights and not to remove outliers directly. Investigating outlier detection performance allowed however to examine what kind of trials are weighted down and how good the method is at detecting such trials. Figure 3 shows all the results for ERP simulated with a SNR of 1. Similar results were observed when using a SNR of 2 (supplementary figure 1). First and foremost, in all cases and for up to 40% of outlying trials, classified trials are located in the upper left corner of the ROC space, indicating good performances. When reaching 50% of outliers, the true positive rate falls down to ∼40% and the false positive rate remains below 40%. This is best appreciated by looking at the plots showing perfect control over false positives when data are contaminated with up to 40% of white, alpha, and gamma outliers. In those cases, the Matthew Correlation Coefficients also remain high (>0.6) although not perfect (not =1), indicating some false negatives. Compared with other types of noise, pink noise elicited very different results, with Matthew Correlation Coefficients around 0 indicating chance classification level. Results from amplitude outliers also show Matthew Correlation Coefficients close to 0 with a linear decrease in true positives and a linear increase in false positives as the percentage of outliers increases. This implies that the PCP method did not detect amplitude changes around peaks. These results are simply explained by the principal components being computed over time frames, and outliers with pink noise and weaker or stronger N1 do not affect the temporal profile of the ground truth sufficiently to lead to different eigenvectors (‘directions’) in this dimension when decomposing the covariance matrix, i.e. their temporal profiles do not differ from the ground truth.

**Figure 3.**
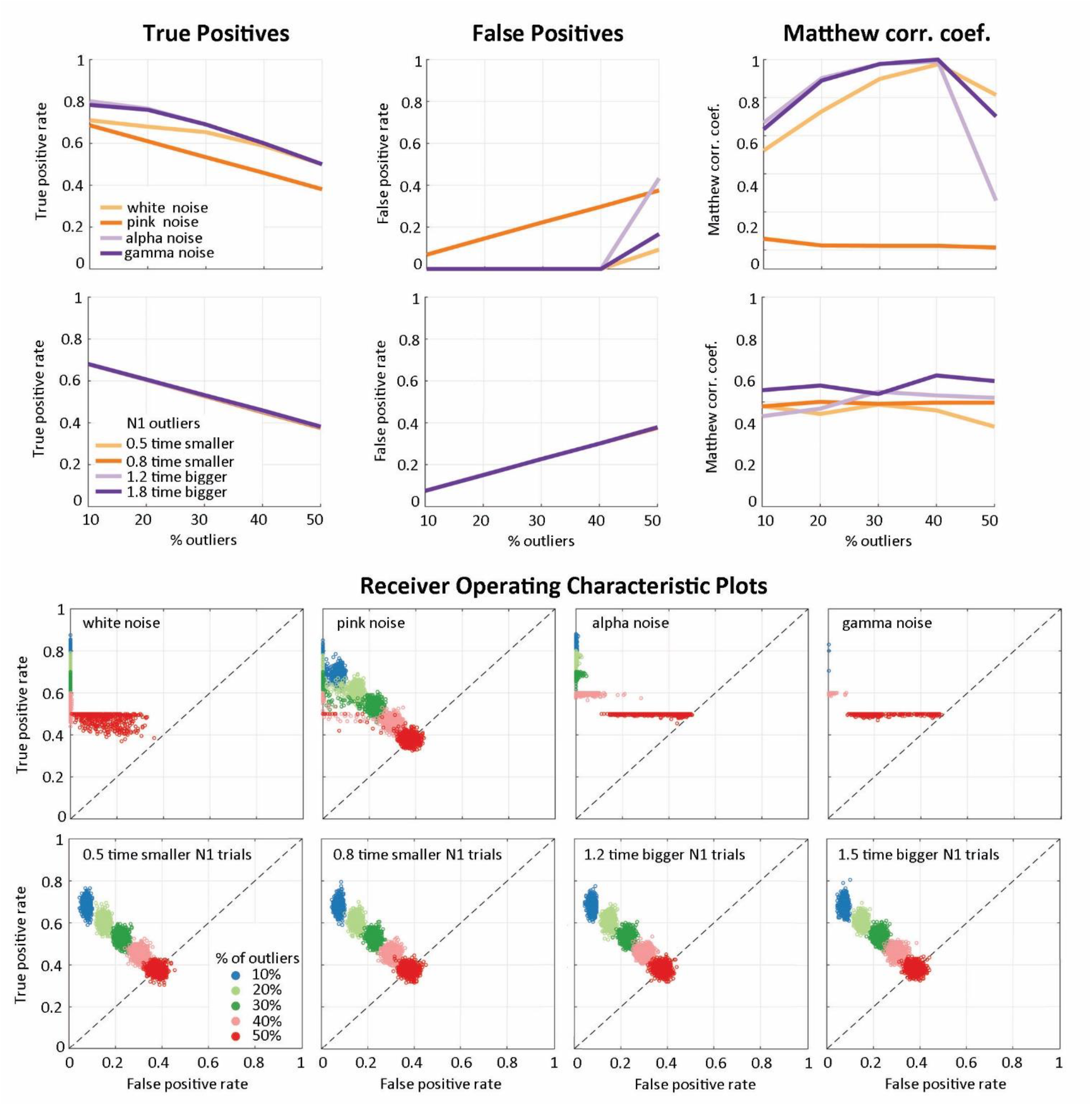
PCP performance at detecting outlying trials with a SNR of 1. (A) Results for outliers affected by white noise, pink noise, alpha, and gamma oscillations. (B) Results for trials affected by amplitude changes over the N1 component (0.5, 0.8, 1.2, 1.5 times the N1). The scatter plots map the Receiver Operating Characteristic Space (False Positive rate vs. True Positive rate); the curves display, from left to right, the median True Positive rate, False Positive rate, and Matthew Correlation Coefficients.

**Supplementary Figure 1.**
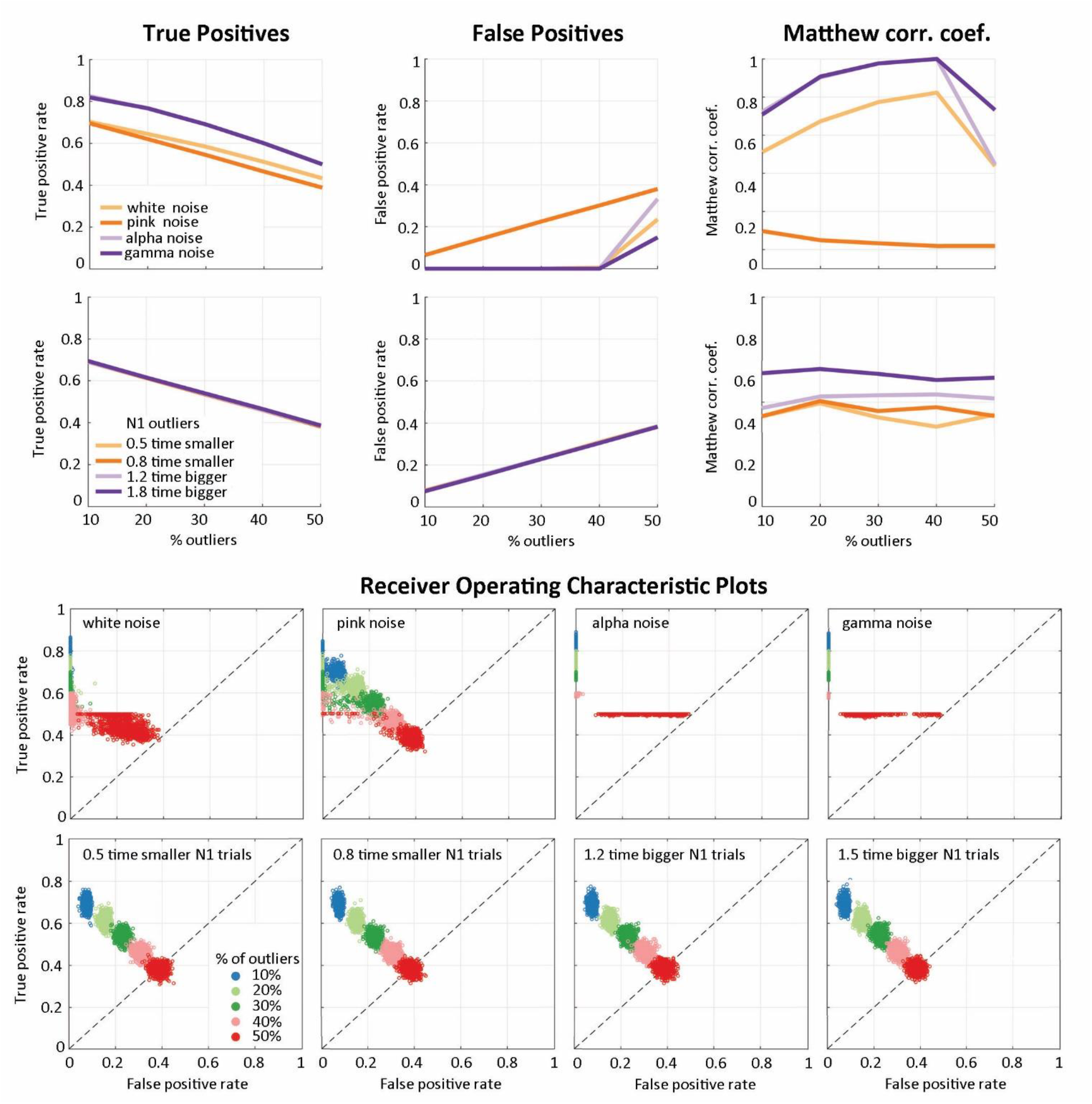
PCP performance at detecting outlying trials with a SNR of 2. (A) Results for outliers affected by white noise, pink noise, alpha, and gamma oscillations. (B) Results for trials affected by amplitude changes over the N1 component (0.5, 0.8, 1.2, 1.5 times the N1). The scatter plots map the Receiver Operating Characteristic Space (False Positive rate vs. True Positive rate); the curves display, from left to right, the median True Positive rate, False Positive rate, and Matthew Correlation Coefficients.

### High vs. low trial weights

The classification for real ERP data confirmed the simulation results: the PCP algorithm weighted down trials with dynamics different from the bulk. Single subject analyses (supplementary table 1) and group analyses (figure 4) for WLS showed that trials with a low weight are less smooth than trials with a high weight (higher temporal variance ∼10 vs. 7.26 μV and power ∼131 vs. 69 dB, lower autocorrelation 11 vs. 12.25 ms), despite having similar spectra (as expected from data filtering and artefact reduction). This is an important result because weights are determined from the multivariate Euclidian distance and skewness of OLS residuals in the Principal Component space, which are independent from those metrics but nevertheless captures those temporal variations.

**Figure 4.**
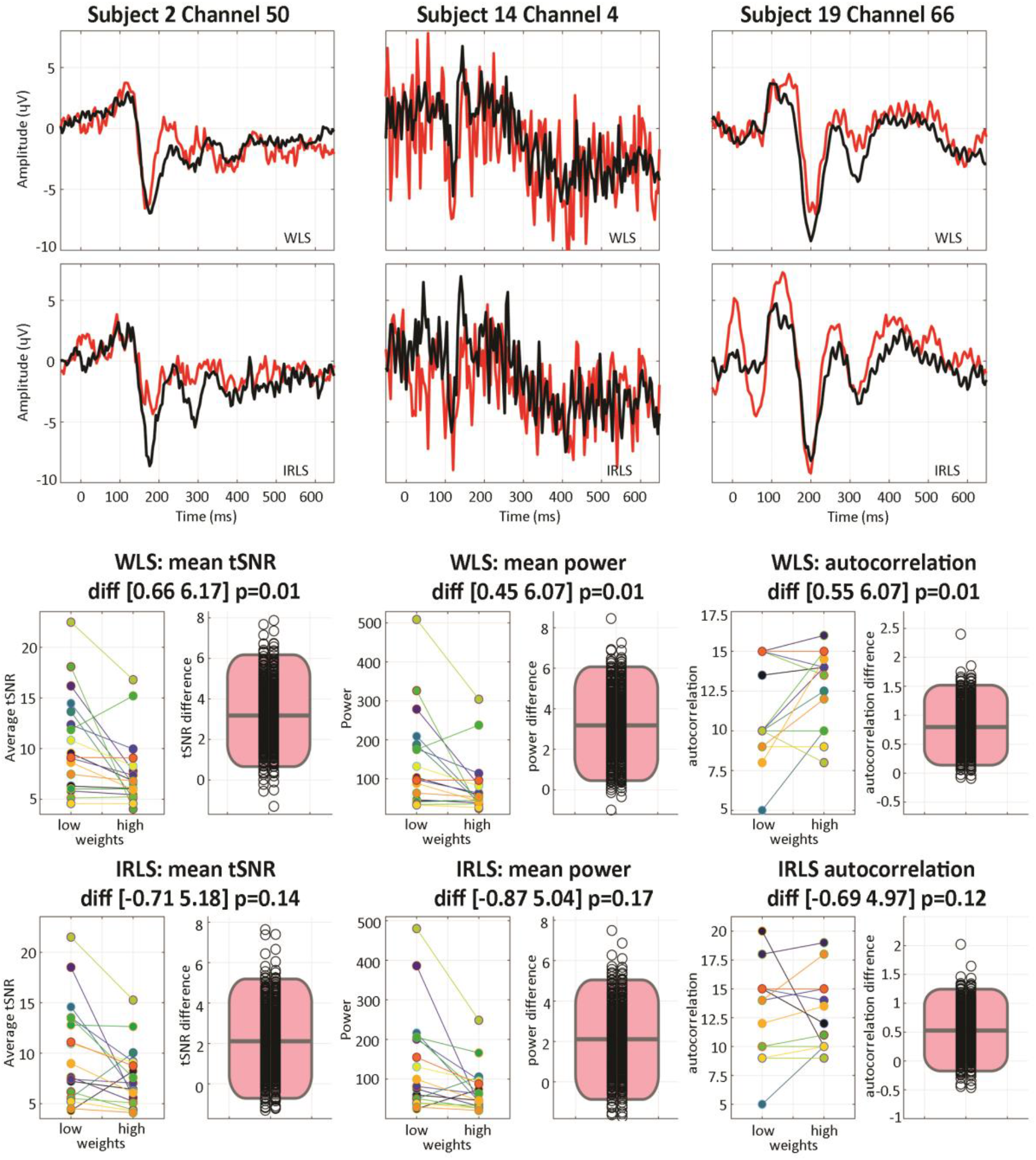
Face ERPs computed for low and high weight trials derived from the Weighted (WLS) and Iterative Reweighted (IRLS) Least Squares approaches. The top of the figure displays the mean of low weight (red) and high weight (black) trials over right posterior temporal (subject 2, channel 50), left frontal (subject 14 channel 4), and left posterior central (subject 19, channel 66) areas showing that in both cases, low and high trials have different dynamics, but also that WLS and IRLS have different weighting resulting in different low and high weight ERPs. The lower part of the figure displays single subject mean tSNR, power and autocorrelation (scatter plots) along with the percentile bootstrap difference between low and high weight trials (black circles are the bootstrap 20% trimmed mean differences and the pink rectangles show the 20% trimmed mean and 95% confidence intervals) revealing that statistical outliers identified by the PCP algorithm (i.e. OLS residual trials with a large distance in the multivariate space) differ in their tSNR, power and autocorrelation, while this is not the case of IRSL outliers.

**Supplementary Table 1.** Subjects 95% percentile bootstrap confidence intervals of 20% trimmed mean differences between high and low trials obtained using PCP-WLS or IRLS at channels with the highest between-trial variance. Intervals which do not include 0 (i.e., the difference between high vs. low trials is statistically significant) are shown on a gray background.

|  | tSNR difference (uV) |  | Power difference (dB) |  | autocorrelation difference (ms) |  |
| --- | --- | --- | --- | --- | --- | --- |
|  | WLS | IRLS | WLS | IRLS | WLS | IRLS |
| s2 | [-0.03 0.54] | [0.26 1.14] | [-2 6] | [3 18] | [-8.5 1.8] | [5.09 16.4] |
| s3 | [2.35 2.92] | [-4.48 -2.34] | [35 50] | [-55 -22] | [-3.9 3.5] | [16.6 45.9] |
| s4 | [0.14 0.69] | [1.9 3.43] | [1 13] | [39 64] | [-13 -6.7] | [-12.8 3.2] |
| s5 | [4.03 8.25] | [10.7 13.57] | [77 200] | [297 382] | [-13 -4.7] | [-14.6 -4.9] |
| s6 | [1.51 2.87] | [-0.74 1.98] | [24 48] | [-6 33] | [-4.8 -0.39] | [-0.6 17.8] |
| s7 | [1.16 5.1] | [2.44 5.26] | [38 141] | [54 129] | [-4 11.1] | [-7.3 11.2] |
| s8 | [7.49 8.21] | [7.57 8.55] | [154 173] | [159 183] | [-24 -19.8] | [-20.2 -14.1] |
| s9 | [2.97 7.96] | [-4.55 0.44] | [52 169] | [-74 28] | [-16 -7.1] | [-1.5 7.1] |
| s10 | [-0.61 0.9] | [-3.47 2.27] | [-11 11] | [-107 102] | [0.9 9.1] | [-0.2 1.5] |
| s11 | [-0.73 4.46] | [4.57 7.27] | [-11 168] | [123 200] | [-2.9 1.4] | [0 7.8] |
| s12 | [6.69 11.17] | [-2.06 4.85] | [149 250] | [-98 93] | [-31 -22] | [-13.1 -2.7] |
| s13 | [-5.06 0.1] | [-6.8 2.91] | [-222 2] | [-285 142] | [4.4 12] | [-6.2 0.19] |
| s14 | [4.81 7.63] | [3.54 7.77] | [174 270] | [123 270] | [-0.4 24] | [-6.9 13.3] |
| s15 | [1.69 3.91] | [-0.97 2.06] | [36 93] | [-20 51] | [-6.5 1.1] | [1.8 10.5] |
| s16 | [-6.85 8.4] | [-2.13 13.82] | [-164 300] | [-65 444] | [-8.3 8.7] | [-16 14.1] |
| s17 | [2.34 3.72] | [2.31 4.09] | [34 68] | [45 83] | [-29.4 -15.9] | [-13.8 2.4] |
| s18 | [0.54 1.28] | [-0.64 1.86] | [6 20] | [-3 27] | [-15.7 -2.43] | [-28.8 11.4] |
| s19 | [-0.39 0.71] | [-0.40 0.57] | [-8 16] | [-9 17] | [-6.9 -1.3] | [-7.1 -1.5] |

By comparison, trials with low and high mean weights based on IRLS, did not differ on those metrics (temporal variance ∼9 vs. 7 μV, and power ∼126 vs. 65 dB, autocorrelation 12.25 vs. 12 ms), which is expected since IRLS weights are computed independently at each time frames. While 11 out of 18 subjects show maximum between-trial variance on the same channels for WLS and IRLS, only 28% of low weight trials were the same between the two methods, and 56% of high weight trials. Since different trials have low or even high weights between methods, this further indicates that the weighting scheme from WLS captures well trial dynamics, differing from IRLS which relies on amplitude variations only.

### Effectiveness of the weighting scheme

The effect of adding outliers on the mean can be seen in figure 5 and supplementary figure 2 in which contaminated data are compared to the ground truth. The standard mean, i.e. the ordinary least squares ERPs, shows an almost linear decrease in Pearson correlations and linear increase in KS distances to the ground truth as the percentage of outlier increases, an expected behaviour since OLS are not robust. Our reference robust approach, IRLS, shows robustness to white noise, alpha, and gamma oscillations with higher Pearson correlations than the OLS. Yet it performed worse than the OLS with pink noise and amplitude outliers, showing lower correlations with the ground truth, despite having similar KS distances in all cases. As the IRLS solution weights data to minimize residuals at each time point separately, these are also expected results, giving an average distance (over time) larger than OLS.

**Figure 5.**
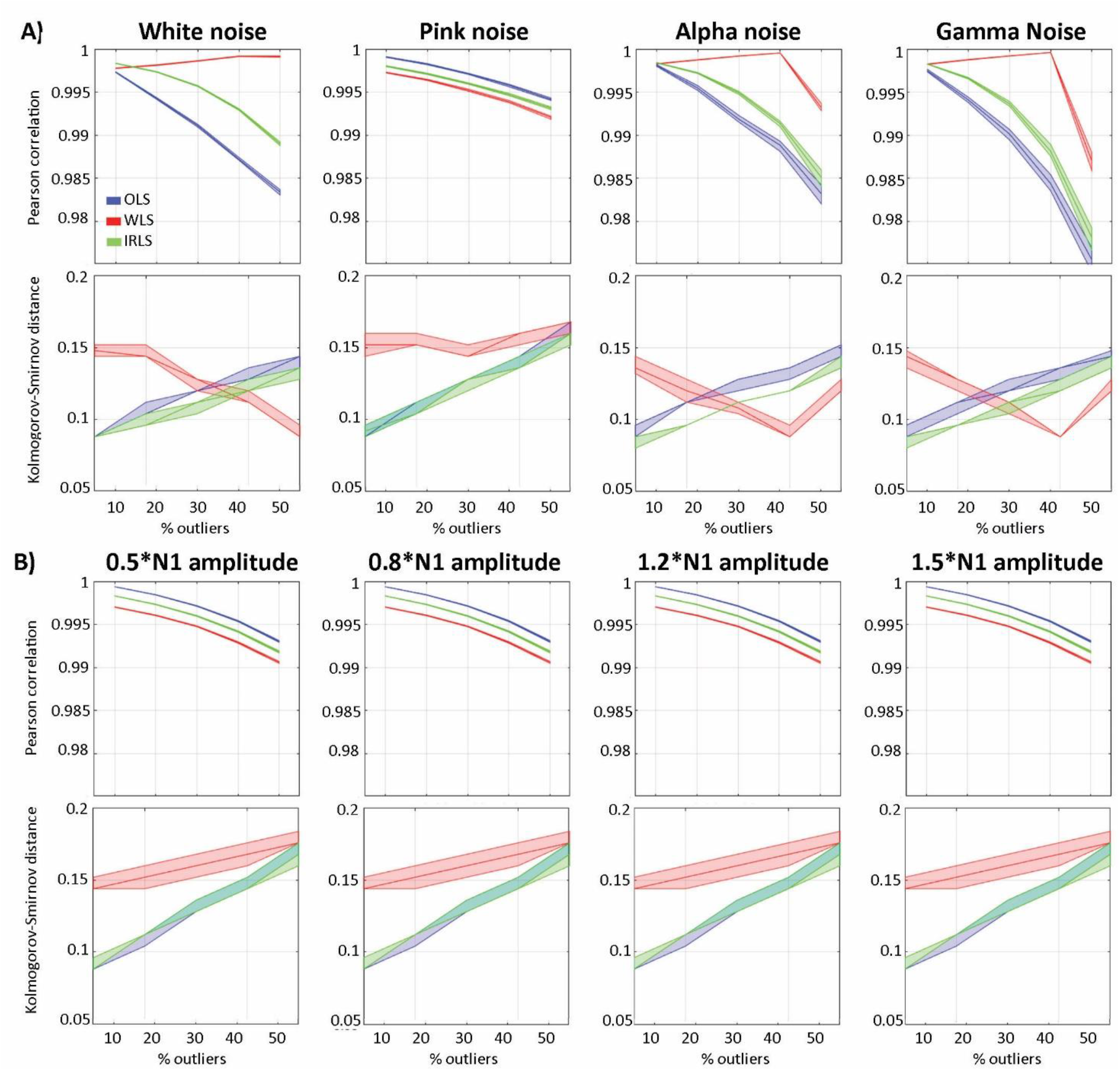
Robustness of the PCP method to outlying trials with a SNR of 1. The upper part of the figure shows median and 95% CI results for outliers affected by white noise, pink noise, alpha and gamma oscillations. The lower part of the figure shows results for trials affected by amplitude changes over the N1 component (0.5, 0.8, 1.2, 1.5 times the N1). Mean Pearson correlations indicate how similar the reconstructed means are to the ground truth (averaged over time), with WLS (in red) largely outperforming OLS (in blue) and IRLS (in green) for white, alpha and gamma noise and higher level of outlying trials, while being less performant for pink noise and N1 amplitude outliers. The mean Kolmogorov-Smirnov distances indicate how much the overall distribution of values differ from the ground truth, showing that WLS ERPs are overall more distant to the (outlier biased) OLS while IRLS still follows the OLS closely.

**Supplementary figure 2.**
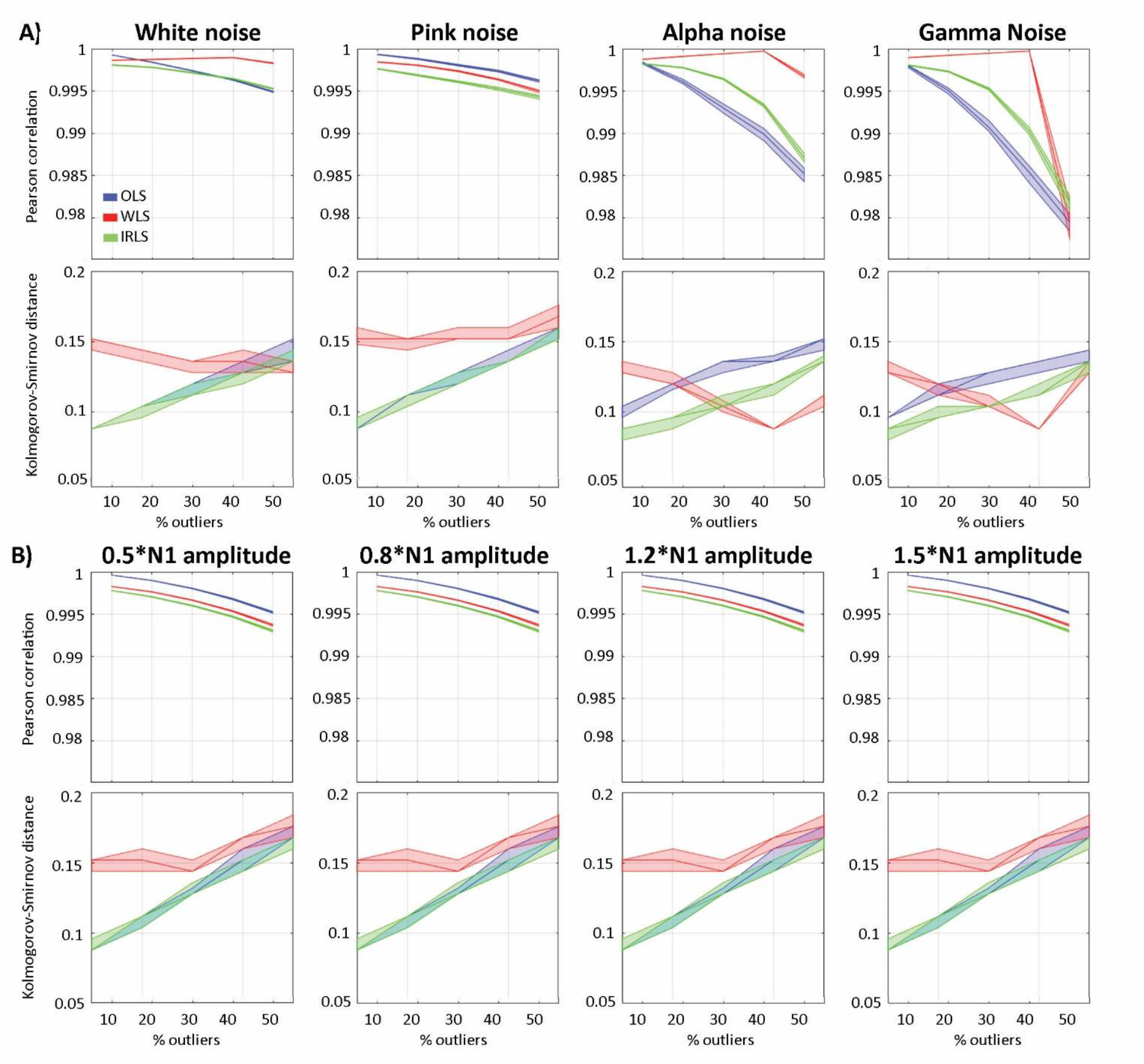
Robustness of the PCP method to outlying trials with a SNR of 2. The upper part of the figure shows median and 95% CI results for outliers affected by white noise, pink noise, alpha and gamma oscillations. The lower part of the figure shows results for trials affected by amplitude changes over the N1 component (0.5, 0.8, 1.2, 1.5 times the N1). Mean Pearson correlations indicate how similar the reconstructed means are to the ground truth (averaged over time), with WLS (in red) largely outperforming OLS (in blue) and IRLS (in green) for white, alpha and gamma noise and higher level of outlying trials, while being less performant for pink noise and N1 amplitude outliers than OLS only. The mean Kolmogorov-Smirnov distances indicate how much the overall distribution of values differ from the ground truth, showing that WLS ERPs are overall more distant to the (outlier biased) OLS while IRLS still follows the OLS closely.

The new WLS approach showed stronger resistance to outliers than OLS and IRLS for white noise, alpha and gamma oscillations, with higher Pearson correlations. For pink noise and N1 amplitude outliers, it performs as well as the IRLS, despite different KS distances (slightly lower correlations with the ground truth for SNR of 1 and slightly higher correlation for SNR of 2). The IRLS algorithm attenuates the influence of those data points that differ from the ground truth, but this may be from different trials at different time points. By doing so, KS distances to the ground truth were similar or lower (for alpha and gamma oscillations) than the OLS. The WLS approach attenuates the influence of trials with different time courses and thus, the WLS ERP mean is affected at every time point, even if the detection concerns a small part of the time course, leading to higher KS distances even with a small number of outliers.

To summarize, simulations showed that the PCP algorithm detects well trials with different dynamics from the bulk leading to WLS-ERP which are, in general, more similar to the ground truth than ERP derived from OLS or IRLS (higher Pearson correlations). Additional results from real-world data indicates that trials with low PCP-WLS weights are trials with different temporal profiles. concurring with the higher Kolmogorov-Smirnov distances observed in the simulation.

### Statistical inference for single subjects

The average type 1 error rate for every channel and time frame tested with simulated data is at the nominal level (5%) for OLS. Results also show that IRLS are a little lenient, with small but significantly smaller p-values than expected, leading to an error rate of ∼0.055. Conversely, WLS are conservative for simulated ERP, with p-values slightly too high, giving a type 1 error rate of ∼0.04) and lenient with purely Gaussian data (type 1 error ∼0.065 – table 1). This behaviour of WLS is caused by the PCP method which optimizes weights based on distances across time, except that with simulated Gaussian data there is no autocorrelation and the PCA returns a much higher number of dimensions, leading to a meaningless feature reduction and thus meaningless trial distances and weights.

**Table 1.** Type I error rate binomial 95% confidence intervals at every time frames and channels for simulated data under the null hypothesis.

|  |  | Null Gaussian | Null ERP |
| --- | --- | --- | --- |
| Regression | OLS | [0.0495 0.0503] | [0.0498 0.0507] |
|  | WLS | [0.0636 0.0645] | [0.0400 0.0408] |
|  | IRLS | [0.0555 0.0564] | [0.0527 0.0536] |
| ANOVA | OLS | [0.0493 0.0502] | [0.0493 0.0501] |
|  | WLS | [0.0695 0.0706] | [0.0374 0.0382] |
|  | IRLS | [0.0575 0.0584] | [0.0540 0.0549] |
| ANCOVA condition | OLS | [0.0494 0.0502] | [0.0493 0.0502] |
|  | WLS | [0.0699 0.0709] | [0.0379 0.0386] |
|  | IRLS | [0.0578 0.0587] | [0.0546 0.0555] |
| ANCOVA covariate | OLS | [0.0496 0.0505] | [0.0496 0.0504 ] |
|  | WLS | [0.0638 0.0648] | [0.0410 0.0418] |
|  | IRLS | [0.0563 0.0572] | [0.0538 0.0547] |

The WLS family-wise type 1 error rate (i.e. controlling the error for statistical testing across the whole data space) examined using nullified ERP data from Wakeman and Henson (2015) shows a good probability coverage for both maximum and cluster statistics with 95% confidence intervals overlapping with the expected nominal value (figure 6). Individual mean values ranged from 0.039 to 0.070 for maximum statistics (across subject average 0.052) and 0.044 to 0.07 for spatial-temporal clustering (across subject average 0.051). Those results do not differ significantly from OLS results (paired bootstrap t-test). Additional analyses based on the number of bootstraps used to build the null distribution indicate that 800 to a 1000 bootstrap samples are enough to obtain stable results, and that the errors are relatively well distributed in space and time even if some channels tend to be more affected than others, i.e. there is no strong sampling bias: maximum number of error occurring at the same location was 0.05% using maximum statistics and 0.9% using spatial-temporal clustering, see bottom for figure 6, error density maps.

**Figure 6.**
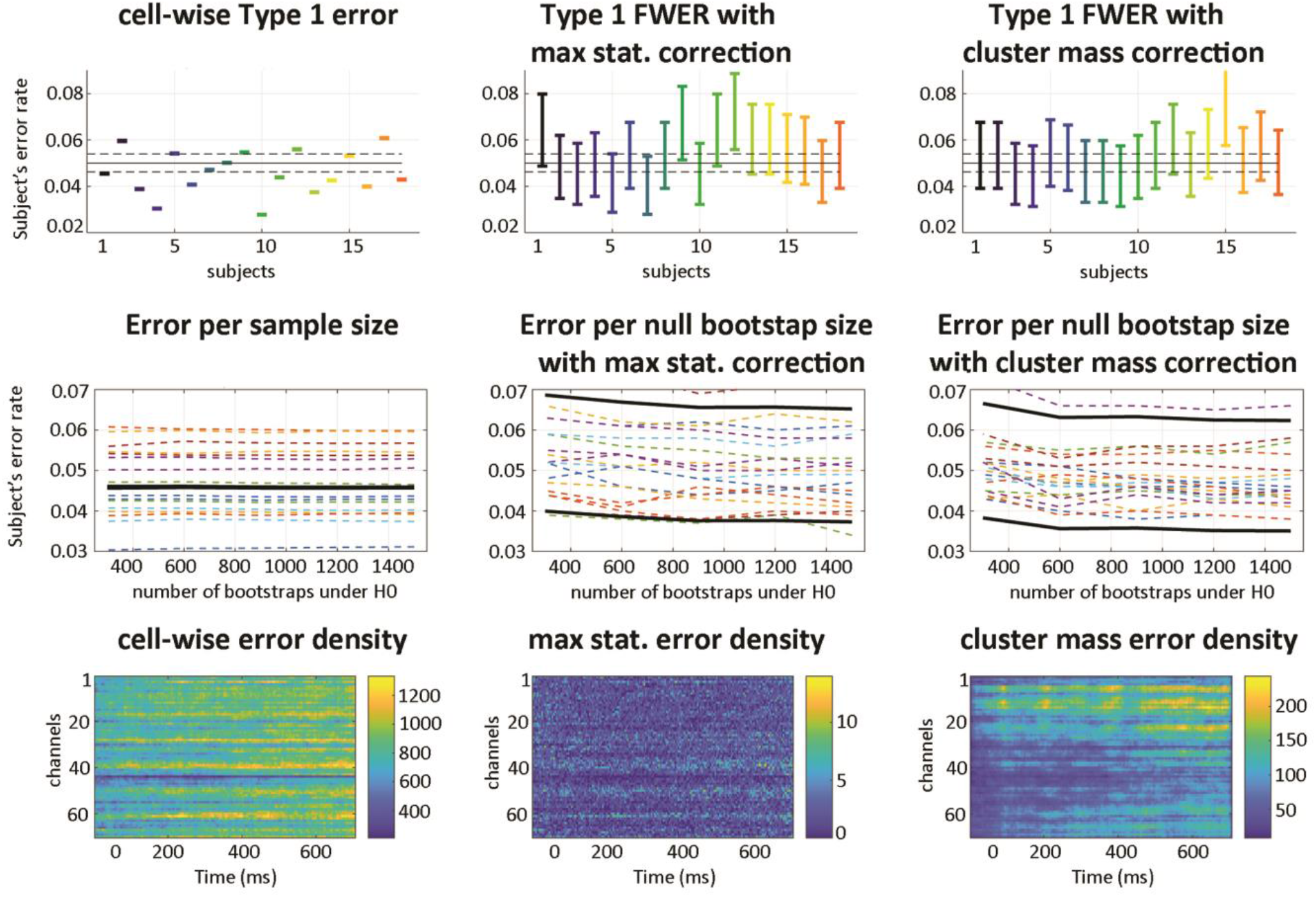
Type 1 error rates under the null using the PCP-WLS method. The top row shows the subjects’ error rates: cell-wise, i.e. averaged across all time frames and channels, and corrected for the whole data space, i.e. type 1 family wise error rate using either the distribution of maxima or the distribution of the biggest cluster-masses. Results are within the expected range (marked by dotted black lines) with overlapping 95% confidence intervals for maximum statistics and spatial-temporal clustering. The middle row shows the effect of the number of resamples, with the dashed lines representing the boundaries of the individual 95% average confidence intervals, and the black lines the average. The cell-wise error is not affected by the number of bootstrap samples since it does not depend directly on this parameter to estimate the null (left). Using maximum statistics and cluster-mass distribution estimates shows a stronger dependency on the number of bootstrap estimates, with results stable after 800 to 1000 bootstraps. The bottom row shows error density maps (sum of errors out of 27000 null maps). The cell-wise error (i.e. no correction for multiple comparisons) shows that errors accumulate, with some channels showing many consecutive time frames with 5% error. By contrast, maximum statistics (middle) and the maximum cluster-masses (right) do not show this effect (maxima at 0.05% and 0.9%), suggesting little to no spatial bias in sampling (note the very different density scales for the three measures).

To summarize, simulations with synthetic data and nullified real-world ERP show that using a Satterthwaite approximation of the degrees of freedom of the error in the context of a WLS solution to the GLM with a single weight per trial, controls well the type 1 error rate.

### Performance evaluation at the group level

Repeated measures ANOVAs using parameter estimates from each method revealed 2 spatial-temporal clusters for the face effect for both WLS and IRLS, but only the 1st cluster was declared statistically significant using OLS (table 2). The expected results (9) with full faces having stronger N170 responses than scrambled faces are replicated for all approaches (start of cluster 1). Maximum differences were observed over the N170 only when using OLS parameters. Using WLS and IRLS gave maxima much later (P280), a result also observed when using TFCE rather than spatial-temporal clustering. In each case, a repetition effect was also observed in a much more consistent way among methods with the second presentation of stimuli differing from the 1st and 3rd presentations (figure 7).

**Table 2:** Face and repetition effects results using cluster-mass correction and TFCE for each of the three methods.

|  | OLS | WLS | IRLS |
| --- | --- | --- | --- |
| Face effect |  |  |  |
| cluster 1 | 140ms to 504ms,<br>max=74, p=0.002 at<br>184ms channel<br>EEG049 | 140ms to 424ms,<br>max=64, p= 0.002 at<br>280ms channel<br>EEG017 | 136ms to 432ms,<br>max=74, p= 0.002 at<br>292ms channel<br>EEG006 |
| cluster 2 |  | 440ms to 648ms,<br>max=17.6, p= 0.032 at<br>616ms channel<br>EEG057 | 520ms to 648ms,<br>max=22, p= 0.032 at<br>636ms channel<br>EEG055 |
| TFCE | max=74, p=0.026<br>at 184ms channel<br>EEG049 | max=64, p=0.012<br>at 280ms channel<br>EEG017 | max=74, p=0.012<br>at 292ms channel<br>EEG006 |
| Repetition effect |  |  |  |
| cluster 1 | 232ms to 648ms,<br>max=50, p= 0.001 at<br>588ms channel<br>EEG057 | 232ms to 648ms,<br>max=51, p= 0.001 at<br>612ms channel<br>EEG045 | 236ms to 648ms,<br>max=52, p= 0.001 at<br>588ms channel<br>EEG057 |
| TFCE | max=50, p=0.002<br>at 588ms channel<br>EEG057 | max=51, p= 0.001 at<br>612ms channel<br>EEG045 | max=52, p= 0.001 at<br>588ms channel<br>EEG057 |

**Figure 7.**
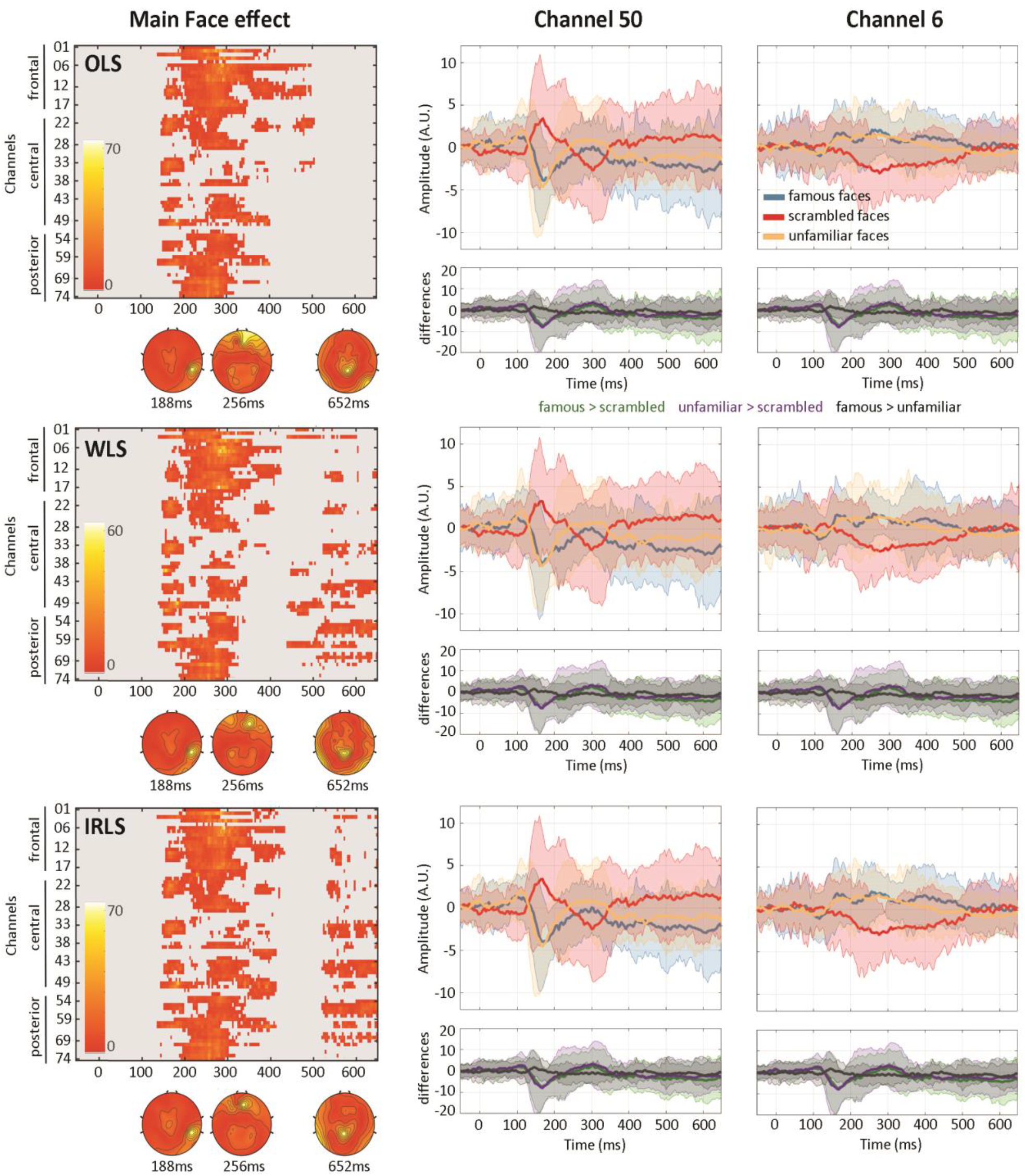
Main Face effects observed using OLS, WLS or IRLS 1st level derived parameters. The left column shows the full channels * times thresholded maps using cluster-mass correction for multiple comparisons (p<.05). Topographies are plotted at three local maxima. WLS and IRLS maps show late effects absent with the OLS solution. The middle and right columns show time courses of the mean parameter estimates per condition (blue, red, orange) and condition differences (green, purple, black) over channel 50 (right inferior-temporal) and channel 6 (middle anterior frontal).

The statistical maps show that group results using based on WLS parameter estimates lead to smaller F values than those obtained from OLS or IRLS estimates (note the difference in maxima table 1 and scale in Figure 7), which is confirmed by the median differences in Hotelling T^2 values (table 3). As expected from a statistical method, the overall shape of ERPs and topographies are unchanged – only effect sizes and significance are affected (supplementary tables 2, 3).

**Table 3.** Median differences in Hotteling T^2 values for each effect tested with percentile bootstrap 95% confidence intervals (p=0.001).

|  | face effect | repetition effect | interaction effect |
| --- | --- | --- | --- |
| WLS vs OLS | -0.32 [-0.36 -0.28] | -0.54 [-0.59 -0.48] | -0.21 [-0.29 -0.13] |
| WLS vs IRLS | -0.34 [-0.39 -0.30] | -0.53 [-0.58 -0.48] | -0.14 -0.21 -0.08] |

Considering uncorrected p-values, this shift in F-values translates into weaker statistical power for WLS: Face effect OLS = 34% of significant data frames, WLS = 31%, IRLS = 34%, Repetition effect OLS = 39%, WLS = 35%, IRLS = 39%. By contrast, results based on cluster-corrected p-values showed however more statistical power for WLS relative to OLS for the Face effect (OLS 20% WLS 22% IRLS 25% of significant data frames with cluster mass and 3%, 5% 3% of significant data frames with TFCE), and mixed results for the Repetition effect (OLS 31% WLS 28% IRLS 31% of significant data frames with cluster mass and 7%, 8% 7% of significant data frames with TFCE). The comparison of distributions’ shapes by deciles (figure 8) provides some understanding of this phenomena. For the face effect, WLS normalized F values tended to be higher but did not differ significantly from OLS or from IRLS, while TFCE values were significantly larger, from the 2nd decile onward when compared to OLS, and for deciles 2, 3, 4, 7, 8 and 9 compared to IRLS. For the repetition effect, WLS normalized F values were also higher and differed from OLS on deciles 2, 7, 8 and 9 for both F-values and TFCE values while it differed from IRLS on decile 9 only when looking at F-values, and deciles 2, 5, 8 and 9 when looking at TFCE values. Finally, for the interaction effect, WLS did not differ from OLS or IRLS in terms of F-values but had significantly weaker TFCE values than OLS (deciles 1, 3, 6, 7, 8 and 9) and IRLS (all deciles but the 4th).

**Figure 8.**
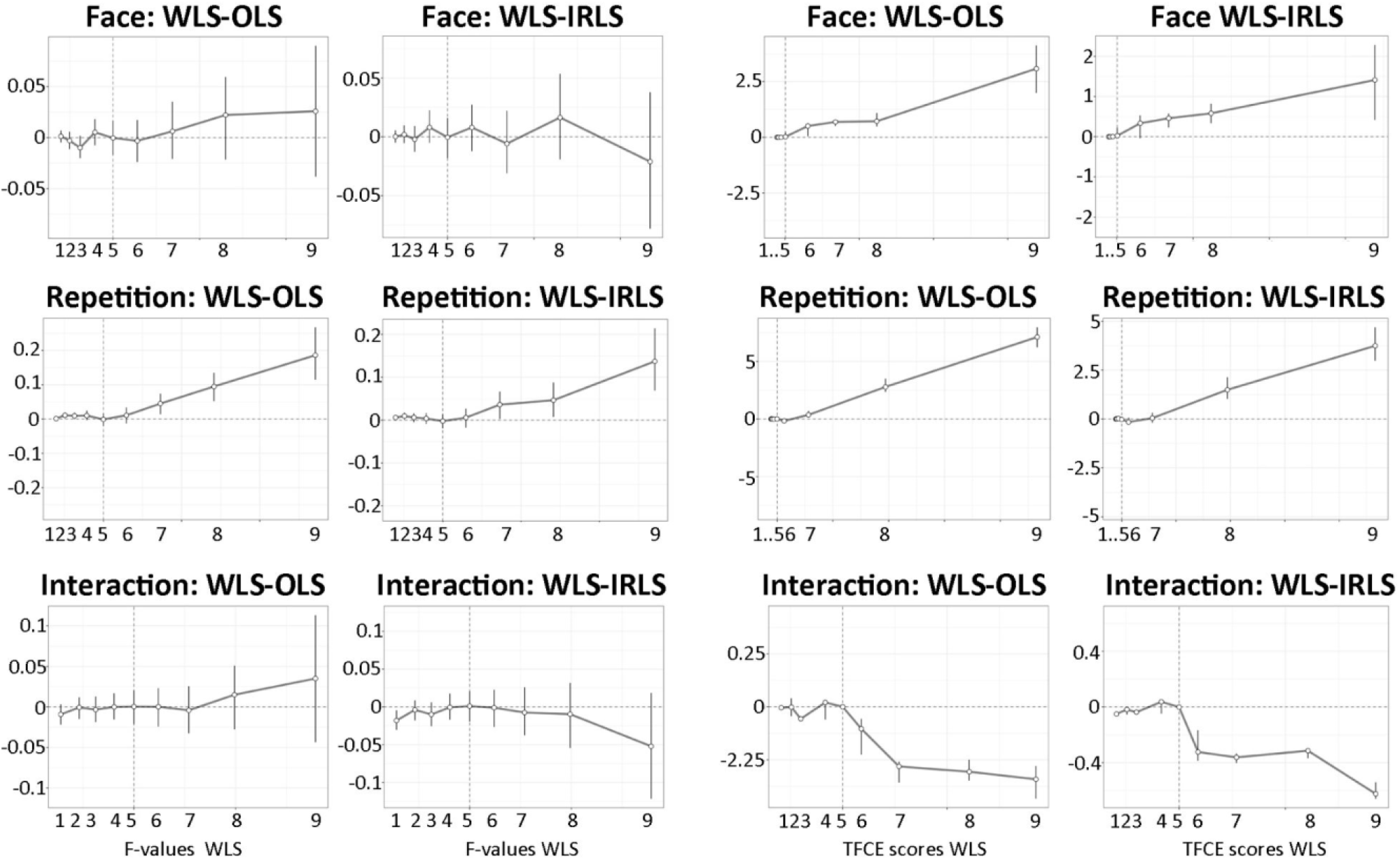
Comparisons of the deciles of standardized F-value (1st and 2nd column) and TFCE value (3rd and 4th column) distributions. Comparisons were done independently for the face effect, the repetition effect and their interaction.

To summarize, for the significant main face effect and main repetition effect, a general pattern of more right skewed distributions of F-values and TFCE-values for WLS than for OLS was observed leading to an increase of statistical power (significance) for cluster corrected analyses, while preserving the topographies (figure 7) and overall ERP shapes (overlap of point estimates with 95% CI of the different methods - Supplementary table 2).

**Supplementary table 2.** Pairwise differences in mean parameter estimates (arbitrary unit) measured at channel 50 and 6 at the maximum of the famous faces responses.

|  |  | OLS | WLS | IRLS |
| --- | --- | --- | --- | --- |
| Cluster 1<br>Channel 50 | Famous Faces vs. Scrambled | -4.93<br>[-12.2 2.32] | -4.52<br>[-11.39 2.34] | -5.82<br>[-12.76 1.11] |
|  | Unfamiliar Faces vs. Scrambled | -4.77<br>[-12.42 2.86] | -4.64<br>[-13.02 3.72] | -5.19<br>[-11.93 1.54] |
|  | Famous vs Unfamiliar Faces | -0.15<br>[-3.13 2.81] | 0.12<br>[-3.28 3.53] | -0.62<br>[-4.86 3.60] |
| Cluster 1<br>Channel 6 | Famous Faces vs. Scrambled | 2<br>[-5.25 9.25] | 1.71<br>[-5.16 8.59] | 1.68<br>[-6.05 9.41] |
|  | Unfamiliar Famous Faces vs. Scrambled | 3.21<br>[-5.80 12.22] | 2.20<br>[-5.97 10.38] | 2.95<br>[-6.08 11.99] |
|  | Famous vs Unfamiliar Faces | -1.20<br>[-5.72 3.30] | -0.49<br>[-5.03 4.04] | -1.27<br>[-5.47 2.93] |
| Cluster 2<br>Channel 50 | Famous Faces vs. Scrambled | -4<br>[-13.82 5.82] | -4.11<br>[-15.62 7.40] | -4.04<br>[-13.31 5.23] |
|  | Unfamiliar Faces vs. Scrambled | -2.16<br>[-9.20 4.87] | -2.17<br>[-9.83 5.48] | -2.32<br>[-8.96 4.31] |
|  | Famous vs Unfamiliar Faces | -1.83<br>[-6.47 2.81] | -1.93<br>[-9.76 5.88] | -1.71<br>[-7.47 4.03] |

**Supplementary table 3.** Medians and maxima of the Hotelling T^2, F-values, Cluster-mass and TFCE scores for each effect of the ANOVA and methods used at the 1st level.

|  |  | medianT | maxT | medianF | maxF | medianCluster | maxCluster | medianTFCE | maxTFCE |
| --- | --- | --- | --- | --- | --- | --- | --- | --- | --- |
| Face effect | OLS | 4.44 | 157.64 | 2.09 | 74.19 | 72.57 | 22591.41 | 130.41 | 40992.1 |
|  | WLS | 3.98 | 136.27 | 1.87 | 64.13 | 64.29 | 19453.52 | 85.72 | 35828.8 |
|  | IRLS | 4.49 | 157.77 | 2.11 | 74.25 | 34.41 | 23300.19 | 130.88 | 54888.48 |
| Repetition Effect | OLS | 5.38 | 107.03 | 2.53 | 50.37 | 35.25 | 39116.91 | 244.38 | 82143.67 |
|  | WLS | 4.46 | 109.14 | 2.1 | 51.36 | 33.76 | 33979.02 | 129.89 | 76244.1 |
|  | IRLS | 5.32 | 110.86 | 2.5 | 52.17 | 37.31 | 39870.66 | 212.27 | 98429.06 |
| Interaction Effect | OLS | 5.45 | 126.31 | 1.12 | 26.01 | 23.79 | 387.94 | 27.64 | 483.46 |
|  | WLS | 5.17 | 78.15 | 1.06 | 16.09 | 21.14 | 317.38 | 25.69 | 470.1 |
|  | IRLS | 5.32 | 135.67 | 1.09 | 27.93 | 30.57 | 283.44 | 22.9 | 366.41 |

## Discussion

Simulation and data-driven results indicate that the proposed WLS-PCP method is efficient at down weighting trials with dynamics differing from the bulk, leading to more accurate estimates. Results show that, for ERP, deriving weights based on the temporal profile provides a robust solution against white noise or uncontrolled oscillations. For biological (pink) noise and amplitude variations which do not alter the temporal profile, the PCP algorithm does not classify well outlier trials, leading to a decrease in detection performance compared with white, alpha or gamma noise. Rather than a defect, we see this as biologically relevant (see below). Importantly, even in those cases of failed detection, the overall correlations with the ground truth remained high (>=0.99). When analyzing real data, differences in amplitude variations were also captured by the PCP/WLS approach, with amplitude variations related to trials which were out of phase with the bulk of the data.

Group-level analyses of the face dataset replicated the main effect of face type (faces>scrambled) in a cluster from ∼150ms to ∼350ms but also revealed a late effect (>500ms), observed when using WLS and IRLS parameter estimates but absent when using OLS parameter estimates. Despite more data frames declared significant with WLS than OLS, effects sizes were smaller for WLS than for OLS and IRLS. The shape of the F distributions when using WLS parameter estimates were however more right skewed than when using OLS or IRLS, leading cluster corrections to declare more data points as significant. Indeed, under the null, very similar distributions of maxima are observed for the three methods leading to more power for the more skewed observed distributions. The interplay between 1st level regularization, 2nd level effect size, and multiple comparison procedures depends on many parameters and it is not entirely clear how statistical power is affected by their combination and requires deeper investigation via simulations. Empirically, we can nevertheless conclude that group results were statistically more powerful using robust approaches at the subject level than when using OLS.

Because the proposed weighing technique relies on a PCA, it is typically requested to have more trials (observations) than data points (time, frequency, or time per frequency), ensuring a full rank covariance matrix. From a design perspective, this is a desirable feature. For a ‘standard’ ERP up to 1 second long, down sampled at 250 Hz, that means 251 trials in total after artefact rejection (about 70 trials per condition for 2 × 2 factorial design). One might argue that is more than usually used, but there is evidence that many more trials than usual are needed for properly powered studies (see (1) for a discussion and references therein). For power analyses, looking at high frequency modulation and thus typically analyzing at 500Hz, this means again 250 data points and thus at least 251 trials, which is more problematic. One can turn to the algorithm to compensate for the lack of trials. One solution may be to use a decomposition for rank reduced data while another may be to down sample the data. Additional simulations varying the number of trials (1500 to 126) and sampling rate (1000Hz, 500Hz, 250Hz) and using white noise outliers, indicate that having more trials than time frames always gives better classification results (as expected) but also that down sampling gives better results than using ill conditioned data (supplementary table 4). Down sampling however interacts with the number of outliers (a priori unknown with real data) such as low sampling rate and high outlier number (>40%) cases have poor performances. In such conditions, computing the data decomposition with ill conditioned data provides better results. As a practical solution, the default usage of the *limo_pcout*.*m* function is thus to down sample the data by 2 if there are not enough trials. If this still results in rank deficient data, the PCP algorithm computes weights on the original ill conditioned data. For spectral analysis in the high frequency domain that will likely suffer from such data conditioning, this still provides a good (automated) solution. It might also be possible to derive better weights (and thus more robust results) by maximizing the multivariate spread of the data in removing more components (i.e. retaining less than 99% of the variance) using e.g. a data driven criteria and/or using a more robust decomposition. This would however requires extensive simulations beyond the scope of this paper which applied the PCP method (4) to create a new WLS scheme.

Using the trial dynamics (temporal or spectral profile) to derive a single weight per trial makes sense, not just because the observed signal is autocorrelated, but also because it is biologically relevant. Let’s consider first the signal plus noise model of ERP generation (20–22). In this conceptualization, ERPs are time-locked additive events running on top of background activity. An outlier time frame for a given trial may occur if 1) the evoked amplitude deviates from the bulk of evoked trials, or 2) the background activity deviates from the rest of the background activity. In the former case, the additional signal may be conceived either as a single process (a chain of neural events at a particular location) or a mixture of processes (multiple, coordinated neural events). In both cases, the data generating process is thought to be evolving over time (auto-regressive) which speaks against flagging or weighting a strong deviation at a particular time frame only. It is likely that several consecutive time frames deviate from most other trials, even though only one time frame is deemed an outlier. In the case of a deviation in background activity, it would mean that for an extremely brief period, a large number of neurons synchronized for non-experimentally related reasons, and for this trial only. Although we do not contend that such events cannot happen in general, this would mean that, in the context of ERP outlier detection, the background activity varies by an amount several folds bigger than the signal, which goes against theory and observations. Let now us consider the phase resetting model (23,24). In this model, ERPs are emerging from phase synchronization among trials, due to stimulus induced phase-resetting of background activity. If a trial deviates from the rest of the trials, this implies that it is out-of-phase. In this scenario, deriving different weights for different time frames (i.e. IRLS solution) means that the time course is seen as an alternation of normal and outlying time frames, which has no meaningful physiological interpretation. Thus, irrespective of the data generating model, the WLS approach seems biologically more appropriate than the IRLS method.

**Supplementary table 4:** additional simulations with white noise and SNR of 1 tested the PCP performance (here Matthew Correlation Coefficients) for different sampling rates and number of trials, in particular having 50% more trials than time frames (1.5 Trial to Frame Ratio (TFR)), just 1 extra (>1 TFR) or just 1 less (<1 TFR) trial or having 50% less trials than time frames (0.5 TFR).

| Sampling frequency | % of outliers | 1.5 TFR | >1 TFR | <1 TFR | 0.5 TFR |
| --- | --- | --- | --- | --- | --- |
| 1000 Hz | 10 % | 0.4248 | 0.4274 | 0.4259 | 0.4379 |
|  | 20 % | 0.5609 | 0.566 | 0.5654 | 0.5815 |
|  | 30 % | 0.6465 | 0.6511 | 0.6509 | 0.6664 |
|  | 40 % | 0.7032 | 0.7079 | 0.7079 | 0.7172 |
|  | 50 % | 0.4238 | 0.3666 | 0.3664 | 0.1835 |
| 500 Hz | 10 % | 0.4307 | 0.4379 | 0.4352 | 0.4555 |
|  | 20 % | 0.5706 | 0.5815 | 0.5804 | 0.6152 |
|  | 30 % | 0.6558 | 0.6664 | 0.6659 | 0.708 |
|  | 40 % | 0.7118 | 0.7172 | 0.7172 | 0.7374 |
|  | 50 % | 0.3049 | 0.1835 | 0.1832 | 0.0593 |
| 250 Hz | 10 % | 0.4417 | 0.4555 | 0.4487 | 0.478 |
|  | 20 % | 0.5937 | 0.6152 | 0.6135 | 0.6671 |
|  | 30 % | 0.6777 | 0.708 | 0.7074 | 0.7806 |
|  | 40 % | 0.7232 | 0.7374 | 0.7375 | 0.2326 |
|  | 50 % | 0.0922 | 0.0593 | 0.0576 | 0.0207 |

In conclusion, we propose a fast and straightforward weighting scheme of single trials for statistical analyses. Results indicate that it captures and attenuates well ERP noise, leading to increased estimation precision and possibly increased statistical power at the group level.

## Acknowledgements

Thank you to EEGLAB/LIMO MEEG users who engaged with the beta version, got stuck but persevered until we solved their issues.

## References

1. Pernet CR, Garrido M, Gramfort A, Maurits N, Michel C, Pang E, et al. Issues and recommendations from the OHBM COBIDAS MEEG committee for reproducible EEG and MEG research. Nature Neuroscience. 2020;

2. Pernet CR, Chauveau N, Gaspar C, Rousselet GA. LIMO EEG: A Toolbox for Hierarchical LInear MOdeling of ElectroEncephaloGraphic Data. Computational Intelligence and Neuroscience. 2011;2011:1–11.

3. Christensen R. Plane Answers to Complex Questions. The theory of Linear Models. 3rd ed. New-York: Springer; 2002.

4. Filzmoser P, Maronna R, Werner M. Outlier identification in high dimensions. Computational Statistics & Data Analysis. 2008;52(3):1694–711.

5. Hoaglin DC, Welsch RE. The Hat Matrix in Regression and ANOVA. The American Statistician. 1978 Feb 1;32(1):17–22.

6. Kasdin NJ. Discrete simulation of colored noise and stochastic processes and 1/f/sup /spl alpha// power law noise generation. Proceedings of the IEEE. 1995 May;83(5):802–27.

7. Yeung N, Bogacz R, Holroyd C, Nieuwenhuis S, Cohen J. Simulated EEG data generator [Internet]. Oxford; 2018. Available from: https://data.mrc.ox.ac.uk/data-set/simulated-eeg-data-generator

8. Satterthwaite FE. An Approximate Distribution of Estimates of Variance Components. Biometrics Bulletin. 1946;2(6):110–4.

9. Wakeman DG, Henson RN. A multi-subject, multi-modal human neuroimaging dataset. Scientific Data. 2015 Jan 20;2:150001.

10. Pernet CR, Martinez-Cancino R, Truong D, Makeig S, Delorme A. From BIDS-Formatted EEG Data to Sensor-Space Group Results: A Fully Reproducible Workflow With EEGLAB and LIMO EEG. Front Neurosci. 2020;14:610388.

11. Pernet CR, Appelhoff S, Gorgolewski KJ, Flandin G, Phillips C, Delorme A, et al. EEG-BIDS, an extension to the brain imaging data structure for electroencephalography. Scientific Data. 2019 Jun 25;6(1):103.

12. Delorme A, Makeig S. EEGLAB: an open source toolbox for analysis of single-trial EEG dynamics including independent component analysis. Journal of Neuroscience Methods. 2004 Mar;134(1):9–21.

13. Onton J, Westerfield M, Townsend J, Makeig S. Imaging human EEG dynamics using independent component analysis. Neuroscience & Biobehavioral Reviews. 2006 Jan 1;30(6):808–22.

14. Pion-Tonachini L, Kreutz-Delgado K, Makeig S. The ICLabel dataset of electroencephalographic (EEG) independent component (IC) features. Data in Brief. 2019 Aug 1;25:104101.

15. Kothe CA, Makeig S. BCILAB: a platform for brain-computer interface development. J Neural Eng. 2013 Oct;10(5):056014.

16. Pernet CR, Latinus M, Nichols TE, Rousselet GA. Cluster-based computational methods for mass univariate analyses of event-related brain potentials/fields: A simulation study. Journal of Neuroscience Methods. 2015 Jul;250:85–93.

17. Maris, E., Oostenveld, R. Nonparametric statistical testing of EEG-and MEG-data. Journal of Neuroscience Methods. 2007;164(1):177–90.

18. Hochberg, Y. A sharper Bonferroni procedure for multiple tests of significance. Biometrika. 1988;75(4):800–2.

19. Rousselet GA, Pernet CR, Wilcox RR. Beyond differences in means: robust graphical methods to compare two groups in neuroscience. European Journal of Neuroscience. 2017;1738–48.

20. Hillyard SA. Electrophysiology of human selective attention. Trends in Neurosciences. 1985;8:400–5.

21. Jervis BW, Nichols MJ, Johnson TE, Allen E, Hudson NR. A Fundamental Investigation of the Composition of Auditory Evoked Potentials. Biomedical Engineering, IEEE Transactions on. 1983 Jan;BME-30(1):43–50.

22. Shah AS. Neural Dynamics and the Fundamental Mechanisms of Event-related Brain Potentials. Cerebral Cortex. 2004 Mar 28;14(5):476–83.

23. Makeig, S., Westerfiled, M., Jung, T-P., Enghoff, S., Townsend, J., Courchesne, E., et al. Dynamic Brain Sources of Visual Evoked Responses. Science. 2002;295(5555):690–4.

24. Sayers BMCA, Beagley HA, Henshall WR. The Mechanism of Auditory Evoked EEG Responses. Nature. 1974 Feb 15;247(5441):481–3.

